# Probing the role of residues lining the active site in the generation of glucose-tolerant variants of a fungal GH1 enzyme

**DOI:** 10.64898/2026.03.09.710506

**Authors:** Barnava Banerjee, Dipayan Chatterjee, Pritha Dasgupta, Chinmay Kamale, Prasenjit Bhaumik

## Abstract

The hydrolytic breakdown of cellobiose into glucose, catalysed by β-glucosidases, is the last and rate-limiting step in cellulose saccharification for producing fermentable glucose in the bioethanol industry. This limitation arises because β-glucosidase activity is inhibited by factors such as temperature, pH, and glucose accumulation in reactors. Enzyme inactivation leads to the buildup of cello-oligosaccharides, which, in turn, inhibit upstream cellulases. Therefore, glucose-tolerant β-glucosidases are preferred for the formulation of industrial cellulase cocktails. In this study, we have recombinantly expressed, purified, and biochemically characterised a β-glucosidase from the cellulolytic fungus *Fusarium odoratissimum* (FoBgl-WT). FoBgl-WT exhibits optimal cellobiose hydrolysis over a broad pH range (4.5-7.5), an important and industrially desirable property for its application in bioreactors. However, the glucose tolerance of FoBgl-WT was ∼0.56 M. Structure-based analyses were carried out to map the residues lining the active site of FoBgl, and their roles in stabilising the product glucose (or even the substrate, cellobiose) were elucidated through a series of site-specific mutations, followed by biochemical characterisation of the resulting FoBgl mutants. Among all the mutants generated, FoBgl-K256I-Y325F exhibits >2.5-fold greater glucose tolerance (∼1.4 M) than FoBgl-WT. Further, we have observed that the FoBgl-K256W and FoBgl-K256I mutants exhibit improved kinetic properties, such as catalytic efficiencies. The structure-based rational engineering efforts improve glucose tolerance and the kinetic properties of FoBgl mutants, making it a useful and promising candidate enzyme for industrial cellulase cocktails.

## Introduction

The tremendous increase in the human population since the Industrial Revolution has been accompanied by a simultaneous rise in energy demand. The energy requirements are met mainly by the consumption of fossil fuels, which have limited reserves (1). The combustion of these fuels harms the environment in various ways, such as the greenhouse effect and environmental pollution (2). In recent years, research into the development of renewable and alternative energy sources has increased. Ethanol is a promising, clean-burning, and renewable alternative to fossil fuels. Ethanol obtained from the hydrolytic breakdown of polysaccharide-rich biomass, followed by fermentation, is referred to as bioethanol. Bioethanol is a promising, clean-burning, and renewable alternative to fossil fuel consumption (3). Bioethanol is being developed for blending with conventional gasoline for automotive fuel, ultimately reducing fossil fuel combustion (4). Depending upon the sources from which bioethanol is generated, it is classified into four generations (5, 6). The first generation of bioethanol is derived from food crops such as sugarcane and maize. This is not a sustainable bioethanol production model, as constant competition exists for using these crops for human consumption and animal fodder (7). The second generation of bioethanol is produced from non-edible carbohydrate-rich sources such as agricultural, municipal, and domestic waste. These materials mainly contain lignocellulose as a significant constituent that can yield fermentable sugars that are ultimately converted to ethanol (7). The third and fourth-generation bioethanol is developed from marine and genetically modified algae, respectively (8). These are still in the initial stages of their development and do not contribute as a significant source of bioethanol due to problems such as excessive water consumption and high costs (9). Thus, among the major sources of bioethanol production, the second-generation bioethanol produced from plant-based lignocellulosic biomass is the most commonly used due to the vast raw material reserves.

Pre-treated and delignified biomass from plant materials is subjected to enzymatic hydrolysis, a process called saccharification, which breaks down long cellulose chains into smaller oligosaccharides (10–14). The synergistic action of cellulase enzymes carries out the enzymatic hydrolysis or saccharification of cellulose (15–17). Cellulases comprise an ensemble of enzymes, including endoglucanases, exoglucanases, and β-glucosidases, which act on cellulose to break it down into monomeric glucose units (18–21). This glucose is fermented into ethanol, which can be used as a fuel. The enzymatic saccharification of cellulose is initiated by endoglucanases, which hydrolyse long chains of cellulose internally. This action of endoglucanases generates more free ends within the cellulose fibre. These free ends are acted upon by exoglucanases, which hydrolyse the chain from these ends to produce cello-oligosaccharides such as cellobiose and cellotriose. β-glucosidases then hydrolyse cellobiose to produce glucose (22, 23). The reaction catalysed by β-glucosidases assumes special interest because it can be considered the rate-limiting step in the cellulose breakdown process. High glucose concentrations inhibit β-glucosidases (24–26). The glucose molecules tend to accumulate in the active-site crater of β-glucosidases and interact with the catalytic residues, forming glucosylated enzymes that are catalytically inactive. Additionally, at high concentrations, glucose molecules displace water from the active site. This inhibits the hydrolytic activity of β-glucosidases and promotes transglycosylation, in which oligosaccharides are generated rather than hydrolysed (27–29). Thus, the inhibition of β-glucosidases leads to the accumulation of higher oligosaccharides in the reactors, which, in turn, inactivates the upstream enzymes in the cellulose saccharification process. Besides higher product concentrations, β-glucosidases are inhibited by higher temperatures and low pH conditions that industrial reactors operate at (29). Hence, there is a need to develop β-glucosidases with enhanced glucose tolerance and optimal hydrolytic activity under the operating conditions of the industrial saccharification process for bioethanol production.

Industrial saccharification of cellulosic biomass is carried out in simultaneous saccharification and fermentation (SSF) reactors, where biomass breakdown and fermentation by yeast occur in the same reactor (30). While this process is economical and has certain advantages, a major limitation is that these reactors operate at 30–35°C, which aligns with yeast’s optimal growth conditions. The cellulases in the cocktail, which are currently used in industry, function at higher temperatures (50-55°C) and exhibit low levels of cellulolytic activity at 30-35°C. Hence, there is a need to explore newer sources of cellulase enzymes that can function at lower temperatures (30-40°C) and be conducive to yeast cell growth. Naturally prevalent β-glucosidases generally do not possess all the qualities required for their applications in the bioreactors for industrial saccharification of cellulose. Hence, protein engineering has emerged over the years as a strong tool for developing enzymes to meet industrial requirements (31). Protein engineering is carried out via two major approaches - directed evolution and rational engineering (32, 33). Directed evolution-based protein engineering has been used to improve the thermal stability of β-glucosidases by introducing random mutations in the DNA sequence. One such example is the engineering of β-glucosidase named Bgl6 obtained from the metagenomic library. Three random mutations were introduced in the DNA sequence coding for *bgl6* to generate a triple mutant M3. The M3 variant of Bgl6 showed improved half-life and thermal stability. However, the M3 mutant possessed poorer glucose tolerance levels than wild-type Bgl6 (34). Another study on a β-glucosidase from *Paenibacillus polymyxa* reported improved half-life and stability through directed evolution. Multiple mutations introduced at or near the active site improved the stability of the enzyme (35). The major disadvantage of directed evolution-based techniques for engineering variants of β-glucosidases with improved glucose tolerance is that random mutagenesis requires a robust screening method to select mutants with desirable properties from a library of mutants. Screening for chemical properties, such as improvements in catalytic efficiency and reductions in product inhibition, has proven more challenging than screening for physical properties, such as thermostability and half-lives. Hence, rational engineering methods have been applied to generate mutants of β-glucosidase to improve these properties. Rational engineering involves the generation of site-specific mutations based on structural information. One study on β-glucosidase from *Trichoderma harzianum* reported enhanced glucose tolerance by reducing the diameter of the catalytic crater opening. This was carried out by substituting smaller amino acids (such as proline or leucine) with bulkier ones (such as tryptophan) to narrow the diameter of the active site lumen (36). Information on the architecture of the catalytic crater, obtained from structural analysis, enabled the identification of specific residues that could be mutated to enhance glucose tolerance in β-glucosidase. Industries have preferred the utilisation of cellulase enzymes from fungal sources owing to their ability to secrete cellulases with high catalytic efficiency (37) in the extracellular environment, and these secreted fungal cellulases naturally function at acidic pH, as cellulolytic fungi grow at this pH.

The current study reports the biochemical characterisation and structure-based rational engineering of a β-glucosidase from the cellulolytic mycelial fungus *Fusarium odoratissimum*, FoBgl-WT. This enzyme was selected for the study because fungal enzymes are expected to exhibit optimal hydrolytic activity at 30-37°C and pH 5.0-6.0, the conditions under which SSF reactors operate. FoBgl-WT has been recombinantly expressed and purified from *Escherichia coli* cells, along with extensive biochemical characterisation. These biochemical assays indicate a unique property of the enzyme, that it retains hydrolytic activity over a broad pH range. Structure-based rational engineering was carried out to improve its glucose tolerance. Based on its structural model, we have designed site-specific mutants with improved glucose tolerance. Our study also sheds light on the importance of the active site architecture and the roles of specific residues lining the active site of FoBgl in influencing its functional properties. Our study would serve as a potential guideline for future protein engineering efforts in improving the functional properties of similar GH1 enzymes.

## Materials and Methods

### Cloning, expression and purification of FoBgl-WT

The codon-optimised gene sequence of FoBgl-WT (NCBI Accession number: XP_031066218.1) for recombinant expression in *E. coli* was synthesised by GeneArt (Thermo Fisher, USA). This gene (*fobgl-wt*) was cloned in the pET28a(+) vector between the *NcoI* and *XhoI* restriction sites. Cloning was performed to insert a hexahistidine tag at the C-terminus of the translated polypeptide. The clones were confirmed by sequencing. Recombinant pET28a(+)*-fobgl-wt* plasmid was isolated from one of the positive clones and used to transform *E. coli* BL21(DE3) cells.

A single colony of transformed *E. coli* BL21(DE3) grown on LA plates with kanamycin (50 µg/ml) was inoculated into LB broth with kanamycin and cultured overnight at 37°C under shaking conditions to serve as the primary inoculum. 1% of this culture was used to inoculate 50 ml LB with kanamycin, grown at 37°C until OD_600_ reached 1.0, induced with 0.4 mM IPTG, and further incubated overnight at 24°C under shaking conditions before harvesting and resuspending the cells in 50 mM Tris-Cl buffer with 400 mM NaCl (pH 7.4). The cells were lysed by sonication, and the soluble and insoluble fractions were separated by centrifugation at 16,000×g for 30 minutes, quantified using the Bradford assay, and analysed by SDS-PAGE.

To obtain FoBgl-WT in bulk amounts, the cells were grown in 1.5 litres of LB, and the same conditions of inducible recombinant expression were used to obtain FoBgl-WT as a recombinant protein in large amounts. The cells were lysed, and the resultant supernatant was loaded on a Ni-NTA column for affinity chromatography (HisTrap FF, 5 ml, Cytiva). The fractions corresponding to purified recombinant FoBgl-WT were pooled, concentrated, and injected into a size exclusion chromatography column - Superdex75 10/300 (Cytiva). All the mutants of FoBgl-WT were expressed and purified in the same manner as described above.

### Biochemical characterisations of FoBgl-WT

Assays were performed with the artificial substrate paranitrophenyl-β-D-glucose (p-NPG) and natural substrate cellobiose for the biochemical characterisations of purified FoBgl-WT. p-NPG assays were carried out by incubating 10 µl of the purified FoBgl-WT enzyme (appropriate concentration) in a reaction mixture comprising 40 µl of 50 mM p-NPG and 450 µl assay buffer at 40°C for 10 minutes. After this, the reaction was stopped by adding 500 µl of 200 mM sodium carbonate. The resultant yellow colour was quantified spectrophotometrically by measuring absorbance at 405 nm. All the assays of FoBgl-WT and its mutants using p-NPG were carried out in the same manner described here. Assays with cellobiose were also carried out to characterise the FoBgl-WT catalysed hydrolysis of the natural substrate. 95 µl of a 350 mM cellobiose solution prepared in a suitable buffer was incubated at 40°C with 5 µl of FoBgl-WT (at appropriate concentrations) for 30 minutes. The reaction was stopped by heating the reaction mixture at 90°C for 10 minutes. The overall glucose concentration was calculated by mixing 2 µl of this reaction mixture with 200 µl of GOD-POD reagent (Accurex) at 37°C for 15 minutes. Using a microplate reader, the resultant pink colour was quantified by measuring the absorbance at 505 nm. All cellobiose hydrolysis assays mentioned in this study have been carried out similarly.

### Estimation of optimum temperature and pH of FoBgl-WT activity

The optimum temperature of p-NPG hydrolysis by purified FoBgl-WT was estimated by preparing the reaction mixture as described previously and carrying out the enzymatic hydrolysis reactions at temperatures of 25°C, 30°C, 37°C, 40°C, 45°C, 50°C and 55°C for 10 minutes. Following this, the reaction was arrested by the addition of Na_2_CO_3._ The optimum temperature of hydrolysis of cellobiose was determined by incubating the reaction mixture as described in the previous section at temperatures of 25°C, 30°C, 37°C, 40°C, 45°C, 50°C and 55°C for 30 minutes.

The optimum pH for p-NPG hydrolysis by FoBgl-WT was determined by carrying out enzymatic reactions in varying pH: 3.5, 4.0, 4.5, 5.0, 5.5, 6.0, 6.5, 7.0, 7.5, and 8.0. 50 mM sodium acetate buffer was used for assays carried out in the pH range of 3.5-5.5, while 50 mM sodium phosphate buffer was used for assays at pH 6.0-7.5, and 50 mM Tris-Cl buffer was used to carry out the reaction at pH 8.0. The p-NPG hydrolysis assay was carried out in the same manner as described previously. The cellobiose hydrolysis assays were performed in the same pH range as described previously. GOD-POD reagent-based assays were performed to quantify the glucose released during the reaction mixture by the action of FoBgl-WT.

All these assays were performed in triplicate. The temperature and pH optima of all the mutants described in this study have been assayed similarly as described above.

### Estimation of glucose tolerance

The glucose tolerance of FoBgl-WT was estimated by carrying out p-NPG hydrolysis assays as described in the section above in the presence of increasing concentrations of glucose. The glucose concentrations used in the assay ranged from 0 - 2 M. The glucose tolerance has been defined as the glucose concentration at which 50% of the enzymatic hydrolysis activity is retained with respect to enzyme activity in the absence of glucose.

The glucose tolerance assays for FoBgl-WT and all its mutants were performed as described above, in triplicate.

### Determination of kinetic parameters

The kinetic parameters were determined for FoBgl-WT and all its mutants by carrying out cellobiose hydrolysis assays. Suitable enzyme concentrations were incubated with increasing concentrations of cellobiose (3.5-332.5 mM) at 40°C for 30 minutes, then the reaction mixture was heated to 90°C for 10 minutes. The overall product yield was estimated using the GOD-POD kit during the process described in the previous section. The data was then fitted into the Michaelis-Menten model using GraphPad Prism 8.4.2, and the kinetic constants of *k_M_*, V_max_, *k_cat_* and *k_cat_/k_M_* were determined.

#### Structural characterisations of FoBgl

For the structural characterisation of FoBgl-WT, the tertiary structure was predicted using the SWISS-MODEL server (http://swissmodel.expasy.org) with GH1 β-glucosidase from *Humicola insolens* (PDB ID: 4MDO) (38) as a template. The model for FoBgl-WT was also generated using the AlphaFold server (39). The two models were superimposed to assess structural differences.

### Molecular dynamics simulations

The generated FoBgl-WT model was simulated for 500 ns to ensure its stability and to obtain energy-minimised sidechain conformations of amino acid residues. The simulations were conducted using GROMACS 2022.2 with the CHARMM36 (40) force field. The starting structure of the protein was prepared using the CHARMM-GUI (41) server. The protein was placed inside a cubic box of water molecules, such that the minimum distance between the protein and the edges of the water box remained almost 7Å. The rest of the steps in preparing the protein structure for MD simulation were identical to those in our previous study (42). It included multiple steps of equilibrating the system using constant temperature, pressure, volume, and number of particles (NVT, NPT). The production MD simulation run was conducted for a 500 ns simulation time, with the atomic coordinates saved after every 10 ps. Following the completion of the simulation time, the important properties such as root-mean-squared deviation (RMSD), root-mean-squared fluctuations (RMSF), radius of gyration (R_g_), distances between atoms of catalytically essential residues, and radial distribution functions (RDF) were calculated. The generated trajectories were visualised using VMD, and the graphs were plotted using GraphPad Prism 8.4.2 software.

#### Site-directed mutagenesis in FoBgl-WT

The residues in the +1 sub-site, selected as probable candidates to affect product binding, included cysteine at the 169^th^ position (C169) and tryptophan at the 168^th^ position (W168). The cysteine residue was mutated to a hydrophobic valine (C169V), and the tryptophan residue was mutated to the aliphatic, hydrophobic leucine (W168L). The +2 sub-site of glucose binding in FoBgl-WT was also analysed. These include the K256 and Y325 residues. Lysine was mutated to isoleucine, leucine and tryptophan (K256I, K256L, K256W, respectively), while tyrosine was mutated to phenylalanine (Y325F).

## Results

### Cloning, expression and purification of FoBgl-WT

The codon-optimised gene coding for *fobgl-wt* was cloned successfully into the pET-28a(+) vector under the control of a T7 promoter with a lac operator to enable inducible expression. Significant amounts of soluble FoBgl-WT could be recombinantly expressed from *E. coli* BL21(DE3) cells by overnight induction of the bacterial culture with 0.4 mM IPTG at 24°C (Figure S1A). This is the first report of recombinant expression of FoBgl-WT.

Once recombinant FoBgl-WT was expressed in sufficient amounts in the soluble fraction, the protein was purified by Ni-NTA affinity chromatography, followed by size-exclusion chromatography. Size-exclusion chromatography yielded FoBgl-WT in highly pure form, as indicated by a single peak. The fractions corresponding to said peaks were analysed on SDS-PAGE gel which yielded a single band corresponding to the molecular weight (55 kDa) of the enzyme (Figure S1B). All the mutants of FoBgl were also purified similarly. The biochemical characterisation of FoBgl-WT and the mutants described in this study was performed using these purified proteins.

#### Biochemical characterizations of FoBgl-WT

##### Optimum temperature, pH for enzymatic activity, and glucose tolerance of FoBgl-WT

FoBgl-WT showed optimal hydrolysis activity at 40-45°C for both p-NPG and cellobiose hydrolysis (Figure 1A). There was a sharp decline in the hydrolytic activity of FoBgl-WT after 45°C, indicating that FoBgl-WT cannot sustain its activity beyond 45°C.

**Figure 1:**
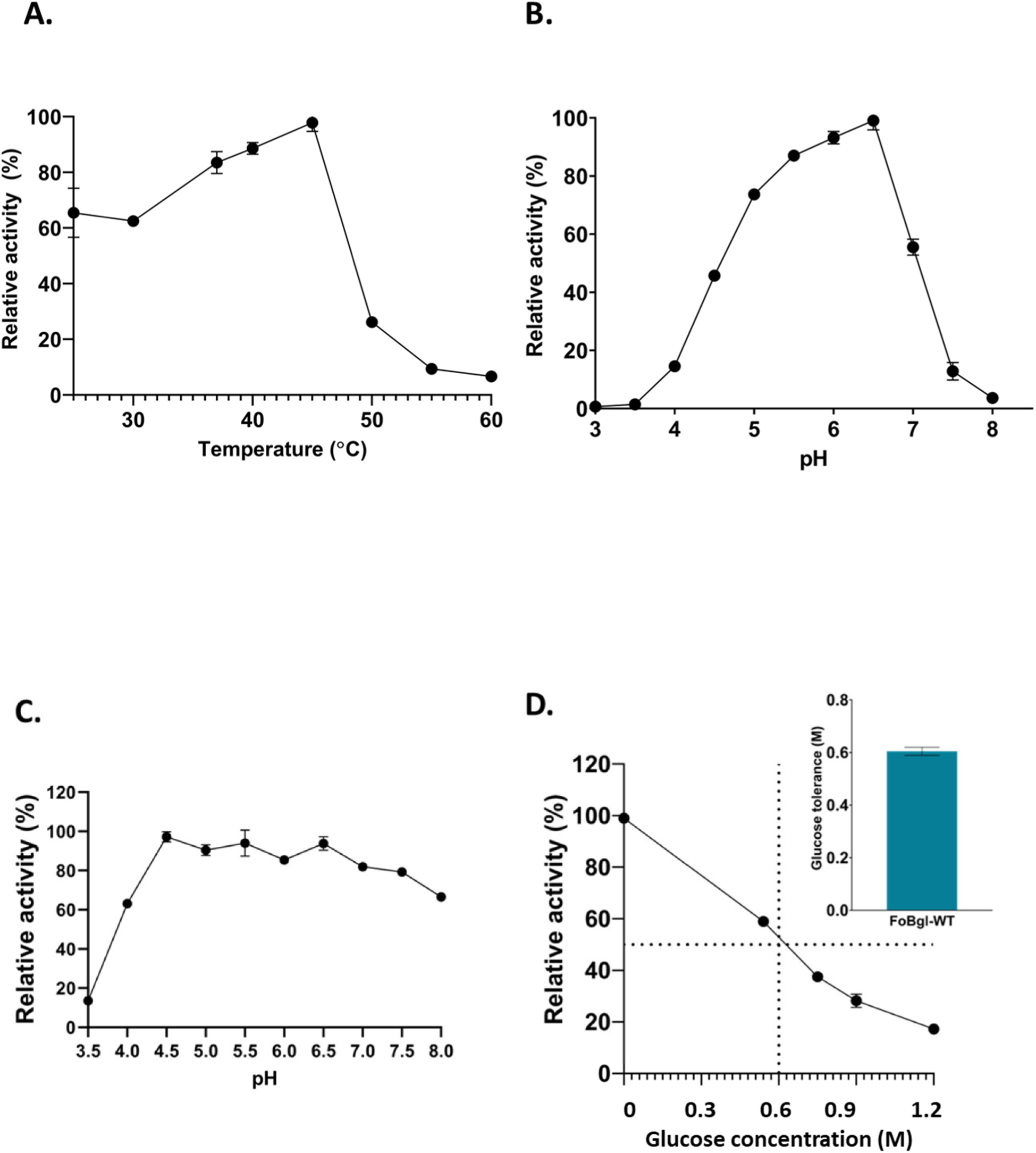
Biochemical characterisations of FoBgl-WT. (A) The optimum temperature of cellobiose hydrolysis. (B) Optimum pH of p-NPG hydrolysis. (C) Optimum pH of cellobiose hydrolysis. (D) Inhibitory effect of glucose on the activity of FoBgl-WT. Inset: Bar graph showing glucose tolerance of FoBgl-WT.

The pH range in which the component enzymes of the cellulase cocktail function is of great industrial importance. Ideally, industrial fermenters operate at a pH of 4.5-5.5; hence, the enzymes that comprise commercial cellulase cocktails should also exhibit optimal hydrolytic activity at this pH range. The pH profile of FoBgl-WT showed an interesting trend. There was a distinct difference between the pH profiles for p-NPG (Figure 1B) and cellobiose (Figure 1C) hydrolysis. The p-NPG hydrolysis profile showed a gradual increase in enzyme activity from pH 3.5 to 6.0, reaching its maximum at pH 6.5. Thereafter, a sharp decline in activity was observed at pH 7-8. The enzyme also showed more than 80% activity at pH 5.0 -5.5, making it a promising candidate for industrial fermenters. However, the cellobiose hydrolysis activity of FoBgl-WT showed a marked difference in the pH profile. There was a gradual increase in enzymatic hydrolysis of cellobiose as the pH increased from 3.5 to 5.0, after which the activity profile remained unaltered, and the enzyme retained >50% activity on cellobiose up to pH 8.0. This is a very promising and unique feature of FoBgl-WT, differentiating it from reported β-glucosidases, as it has hydrolysis activity on cellobiose over a wide pH range, making it a suitable candidate for industrial cellulase cocktails. Industrial cellulases require a glucose tolerance of ∼0.8 M. Recombinantly expressed and purified FoBgl-WT shows a glucose tolerance of ∼0.56 M (Figure 1D). Hence, there is a need to further enhance the glucose tolerance of FoBgl-WT for use in industrial reactors.

##### Kinetic characterisation of FoBgl-WT

The enzyme kinetics assays of FoBgl-WT were performed with cellobiose as a substrate. The kinetic parameters of FoBgl-WT have been summarised in Table 1. As the *k_cat_/k_M_* of enzymatic reactions indicates the catalytic efficiency, the effect of the various mutations performed on the catalytic efficiency of FoBgl has also been documented in this study.

**Table 1:** Kinetic properties of FoBgl-WT and all the mutants that show cellobiose hydrolysis (all the assays were performed in triplicate)

| **Mutants** | **$k_M$ (mM)** | **$V_{max}$**<br>**($\mu\text{mol}/\text{min}/\text{mg}$)** | **$k_{cat}$ ($\text{min}^{-1}$)** | **$k_{cat}/k_M$ ($\text{mM}^{-1}\text{min}^{-1}$)** | **Glucose Tolerance(M)** |
| --- | --- | --- | --- | --- | --- |
| FoBgl-WT | 15.55 ± 3.96 | 9.95 ± 0.51 | 547.25 | 35.19 | 0.56 |
| FoBgl-K256W | 9.71 ± 2.36 | 16.50 ± 0.84 | 680 | 69.99 | 0.9 |
| FoBgl-K256I | 17.81 ± 3.4 | 12.03 ± 0.40 | 661.38 | 37.13 | 1.2 |
| FoBgl-K256L | 59.18 ± 11.51 | 17.35 ± 0.23 | 954.36 | 16.12 | 0.8 |
| FoBgl-Y325F | 11.14 ± 1.18 | 1.59 ± 0.02 | 88.08 | 7.89 | 1.1 |
| FoBgl_K256W-Y325F | 11.97 ± 1.37 | 4.95 ± 0.14 | 11.08 | 0.92 | 1.2 |
| FoBgl-K256I-Y325F | 37.03 ± 4.74 | 6.91 ± 0.22 | 7.95 | 0.21 | 1.4 |

#### Structural analyses of FoBgl-WT reveal potential sites for mutations

In the absence of an experimentally determined crystal structure of FoBgl, the structure of the enzyme was predicted so as to identify the residues along the catalytic crater. The 3D structure was predicted using both homology modelling and AlphaFold. The homology model of FoBgl-WT was built using the SWISS-MODEL server, and it was built from sequence-level information of FoBgl-WT using AlphaFold 2.0. The two structures were superposed (Figure S2A) to analyse their differences. Apart from a stretch of amino acid residues from 475-490, both models were identical (RMSD of the Cαs of the residues of the aligned structures was calculated to be 0.36 Å). The AlphaFold 2.0 predicted an unstructured random coil for the corresponding residues (475–490). The confidence scores of the prediction for these C-terminal residues in the AlphaFold-generated model were low compared to the >90% pLDDT scores for the other regions in the protein structure. Since the two models were almost identical, further analyses were carried out using the structure predicted by SWISS-MODEL. This is because the SWISS-MODEL structure was generated using the well-characterised β-glucosidase from *Humicola insolens* as the template, and we were more confident in the accuracy of the side-chain conformations based on the experimentally determined structure as the template.

The overall structure of FoBgl-WT showed the characteristic (α/β)_8_-barrel fold (Figure 2A), which one of the characteristic features of the β-glucosidases belonging to the GH1 family. The generated FoBgl-WT model was compared with structures of other GH1 family β-glucosidases. Both the sequences and experimentally determined crystal structures of these enzymes were considered for comparison. The active site of FoBgl-WT was found to exist as a deep, blind-ended crater with the catalytic glutamates present at the bottom of this crater. The catalytic glutamate residues are present in two conserved catalytic motifs, TXN**E**P (where X denotes any amino acid) and I/YT**E**NG. β-glucosidases from the GH1 family of enzymes exhibit a retention mechanism of cellobiose hydrolysis, where the stereochemistry of the anomeric carbons in the product glucose molecules is identical to that of the substrate (42). To carry out hydrolysis via the retention mechanism, the delta carbons (CD) of the two catalytic glutamates are expected to be at a distance of ∼5.0-5.5 Å (43). Several published structural and biochemical studies on GH1 family β-glucosidases identify the C-terminal glutamate residue as the catalytic nucleophile, which initiates the nucleophilic attack on the β-(1,4) glycosidic bond. The deprotonated state of the C-terminal glutamate is needed for this nucleophilic attack. The N-terminal glutamate residue, acting as a catalytic acid/base, is involved in the second half of the hydrolysis reaction, in which the glucosylated enzyme intermediate is cleaved by a water molecule activated by proton abstraction from this glutamate residue. To gain a detailed understanding of the enzyme’s inherent dynamics and molecular interactions at the structural level, the FoBgl-WT model was simulated for 500 ns.

**Figure 2:**
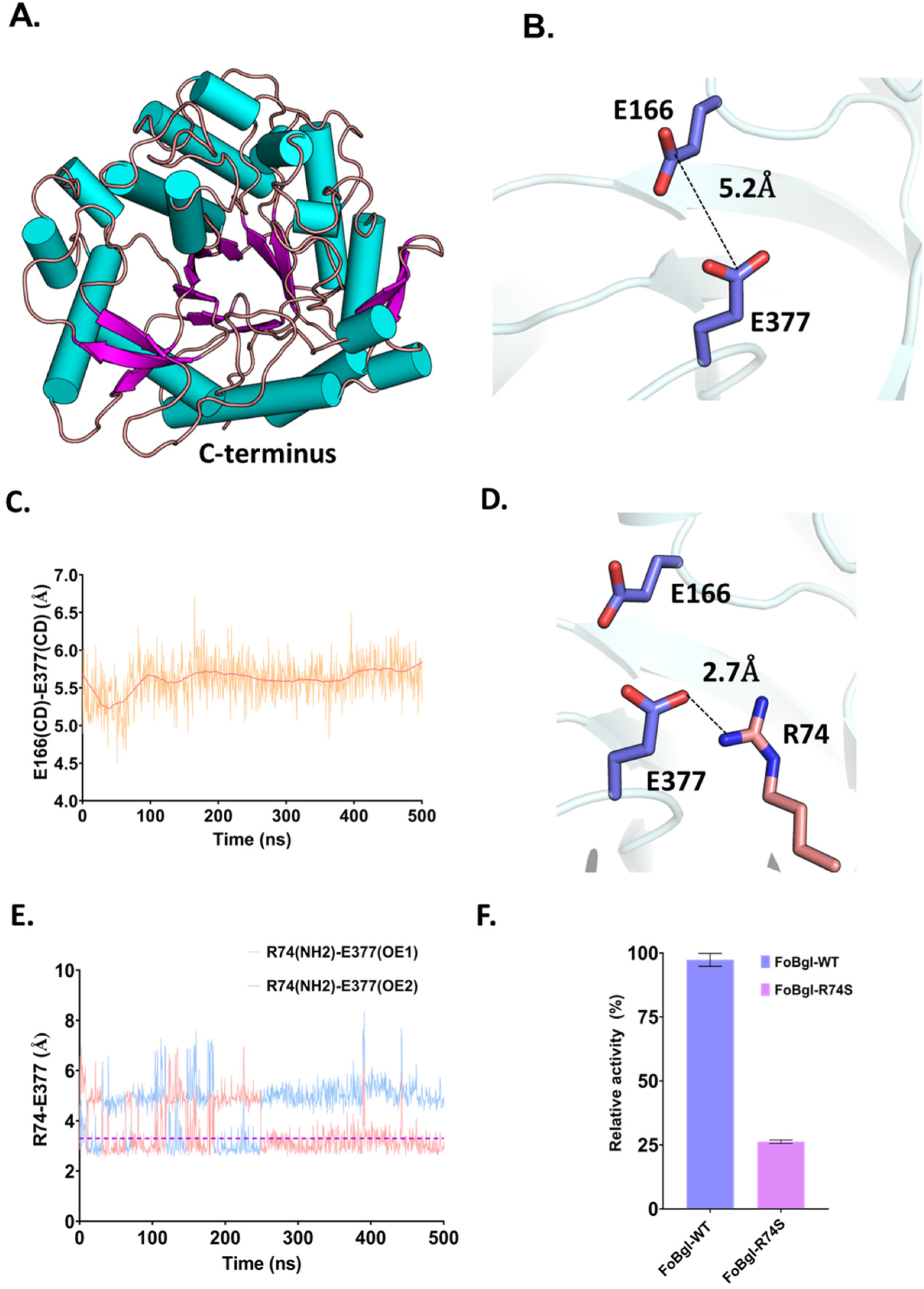
Structural features of FoBgl-WT. (A) The overall structure of FoBgl-WT shows a distinct TIM barrel fold, with secondary structures represented in different colours. (B) Zoomed-in view of the catalytic glutamates (E166 and E377) represented as blue sticks. (C) Distance between the CD of E166 and E377 analysed from the 500 ns MD simulation trajectory. (D) Representation of the R74 and E377 residues in FoBgl showing the distance between the atoms of interest. (E) Maintenance of the salt bridge interaction between E377 and R74 throughout 500 ns of MD simulation. (F) Comparison of cellobiose hydrolysis activity of FoBgl-WT and FoBgl-R74S mutant indicates the importance of the salt bridge interaction.

**Figure 3:**
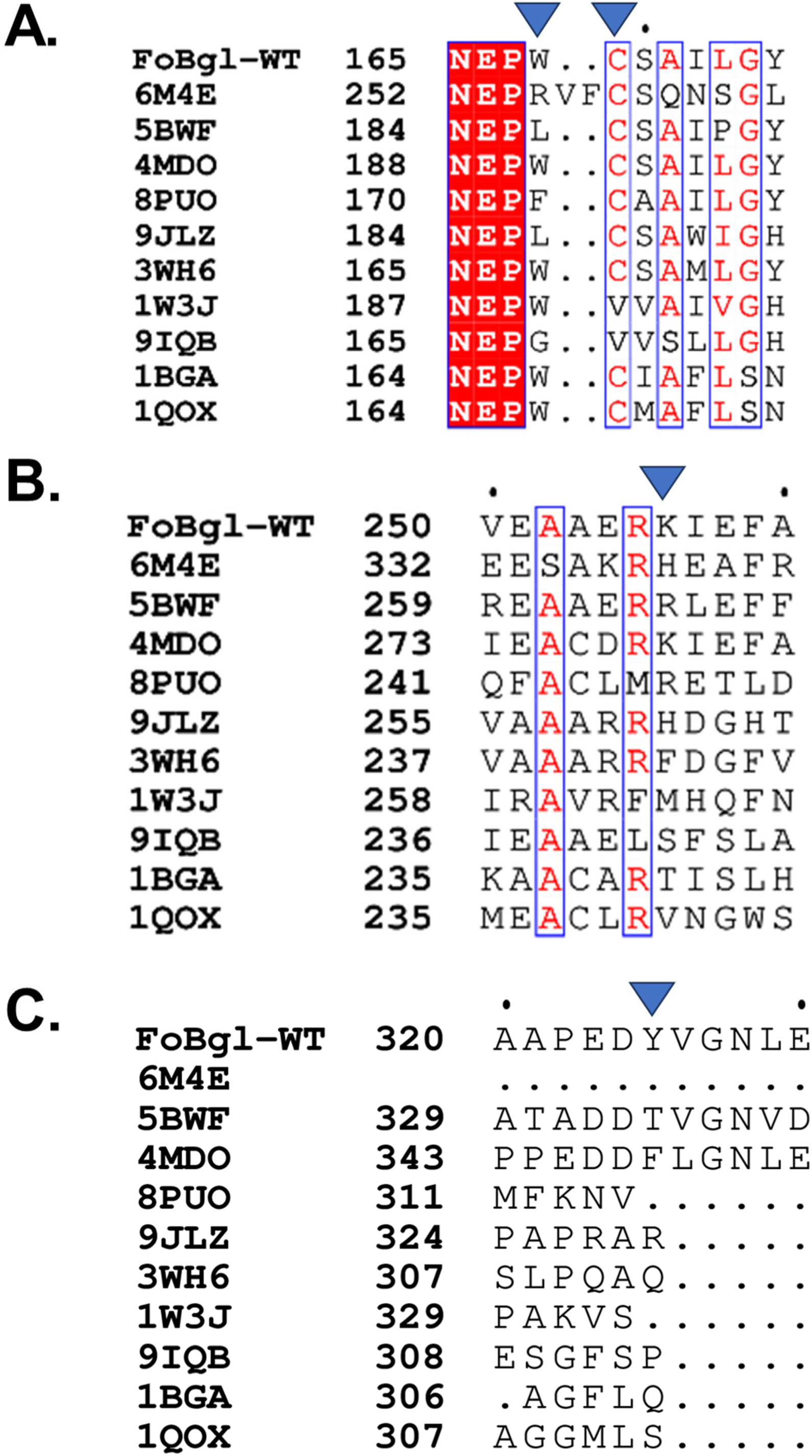
Sequence comparisons of well-characterised GH1 enzymes with FoBgl-WT. (A) Sequence comparisons of +1 sub-site residues of FoBgl-WT with 10 structurally characterised GH1 enzymes. The blue arrows indicate the residues in FoBgl-WT that were selected for site-specific mutagenesis. (B and C) Sequence comparisons of the +2 sub-site residues of FoBgl-WT with 10 structurally characterised GH1 enzymes. PDB IDs of different GH1 enzymes are mentioned as sequence identifiers. 6M4E: enzyme from *Hamamotoa singularis*, 5BWF: enzyme from *Trichoderma herzanium*, 4MDO: enzyme from *Humicola insolens*, 8PUO: enzyme from an Antarctic Marinomonas, 3WH6: enzyme Td2F2 from the metagenome, 1W3J: enzyme from *Thermotoga maritima*, 1BGA: enzyme from *Bacillus polymyxa*, 1QOX: enzyme from *Bacillus circulans,* 9JLZ: enzyme from the soil metagenome (UnBGl1), and 9IQB: enzyme from *Acetivibrio thermocellus*.

### Molecular insights into the active sites of FoBgl to understand the role of its catalytic residues using molecular dynamics simulations

MD simulations were performed to examine the stability and structural dynamics of the FoBgl-WT model under conditions mimicking physiological conditions. The simulations used the FoBgl-WT structure in an aqueous environment, which was generated by placing the protein in a water box of suitable dimensions, with ions, to mimic physiological conditions. The model remained stable for 500 ns of the MD simulation, as indicated by the correlation between RMSD and radius of gyration (Figure S2B). The fluctuations in different residues during the simulations have been recorded using the RMSF analysis (Figure S2C). The greatest degrees of fluctuations have been observed in residues situated near the loop regions of the proteins.

The distance between the two catalytic glutamates remains within 5.0-5.5 Å throughout the time period of the simulation, indicating that the side chain conformations of the catalytic residues depict the catalytically competent state of the enzyme (Figure 2B, C). Apart from distance, the protonation states of the catalytic glutamates, which in turn are a function of the micro-environment, play a very important role in the catalytic process. As previously described, the biochemical studies on FoBgl-WT revealed that the enzyme retains its catalytic activity on cellobiose over a broad pH range. We wanted to investigate the molecular determinants underlying this. E166, the probable residue to serve as the general acid/base in the catalytic event, abstracts a proton from nearby water molecules. On the other hand, the E377 residue, which serves as the C-terminal catalytic glutamate, was observed to form a salt bridge interaction with the R74 residue. The salt bridge was observed to be maintained throughout the 500 ns duration by engaging either the OE1 or the OE2 atoms of the E377 residue (Figure 2D, E). This salt-bridge interaction with a conserved arginine would be essential in maintaining the E377 residue in the deprotonated state, which drives the nucleophilic attack on the glycosidic bond of the sugar substrates. We have further investigated this claim by mutating R74 to serine, which is much smaller and lacks a positive charge. The effect of this substitution was manifested in the catalytic activity of the FoBgl-R74S mutant. This mutant retained only ∼27% of the cellobiose hydrolysis activity of the wild-type enzyme (Figure 2F). This shows that the conserved R74 residue is crucial in maintaining the microenvironment around the catalytic nucleophile E377. Our computational analyses indicate that the protein system used in simulations is catalytically competent, highlighting the role of the microenvironment around the catalytic residues in facilitating reaction progression.

### The catalytic crater of FoBgl-WT

The FoBgl-WT structural model revealed the presence of the catalytic residues located at the base of the catalytic crater, a feature observed in GH1 β-glucosidases. The catalytic crater is lined with several amino acid residues, which are essential in stabilising or binding the substrate or product molecules. The cross-section of the catalytic crater in the FoBgl-WT structure shows residues such as K256, C169, Y325, N306, T233, D237, N311, and N165. These residues might have a role in stabilising sugar moieties through the polar interactions involving the OH group in these sugars (Supplementary Movie M1). Besides these polar interactions, sugars can also be stabilised inside the catalytic crater via stacking interactions (Supplementary Movie M1). These interactions are provided in FoBgl-WT by hydrophobic and aromatic amino acid residues such as W121 and W168. These tryptophan residues can interact with the sugar backbone, thereby causing the sugar moieties to bind within the catalytic crater.

An in-depth structural understanding of the catalytic crater architecture is key to generating site-specific mutations that aid protein engineering. As discussed previously, poor glucose tolerance is a major bottleneck to the application of β-glucosidases in the saccharification process. Enhancement of glucose tolerance can be achieved by altering the residues that stabilise glucose in the catalytic crater. The mutations were designed to prevent the glucose generated as a product of cellobiose hydrolysis from binding in the enzyme’s active site.

#### Site-directed mutagenesis of FoBgl-WT to enhance glucose tolerance

An extensive sequence-based and structural comparison of the FoBgl-WT model with those of well-characterised glucose-tolerant GH1-family β-glucosidases was performed. The main focus was on determining the role of residues lining the catalytic crater contributing to the glucose tolerance and catalytic performance of FoBgl, as we hypothesised that these residues play essential roles in stabilising the substrate or product. The cross-sectional view of the catalytic crater of FoBgl-WT was compared with that of a GH1 β-glucosidase, UnBGl1(44).UnBGl1 is a β-glucosidase obtained from the soil metagenome, and our group has crystallised it in both apo and glucose-bound forms. The glucose-bound form of UnBGl1 shows the presence of three glucose molecules inside the catalytic crater, occupying conserved sites, which have been named -1, +1, and +2 sub-sites, respectively (Figure S3A). The -1 sub-site comprises the two catalytic glutamates (E166 and E77). The +1 sub-site includes residues W168 and C169. These residues can interact with the product glucose via stacking or polar interactions. C169 was mutated to valine, and tryptophan was mutated to leucine to prevent such interactions. Also, at the +2-sub-site forming an entry of the catalytic crater, we observed a lysine 256 residue that could form polar interactions and stabilise glucose (Figure?). This lysine was mutated to tryptophan and aliphatic non-polar residues such as isoleucine and leucine to prevent such interactions. Along with this lysine residue, a tyrosine 325 residue was observed at the dorsal end of the opening of the catalytic crater. This tyrosine was mutated to phenylalanine to prevent the OH group from participating in polar interactions with glucose. All the mutants described above were expressed and purified individually. Their biochemical properties, such as temperature and pH optima for cellobiose, p-NPG hydrolysis, and glucose tolerance, were determined. Furthermore, the kinetic properties of these mutants were determined and compared with FoBgl-WT to understand the molecular determinants of catalytic efficiency and turnover number.

#### Mutations at the +1 sub-site of sugar-binding

W168 and C169 residues were observed to be a part of the +1 sub-site inside the catalytic crater of FoBgl. Structure-based comparisons with the glucose-bound structure of UnBGl1 revealed that both these residues could form polar interactions leading to stabilisation of glucose inside the active site, thereby causing inhibition of FoBgl-WT activity at higher glucose concentrations. W168 was mutated to leucine (W168L), and C169 was mutated to valine (C169V) (Figure 4A). These mutants were expressed and purified in a manner identical to the FoBgl-WT.

**Figure 4:**
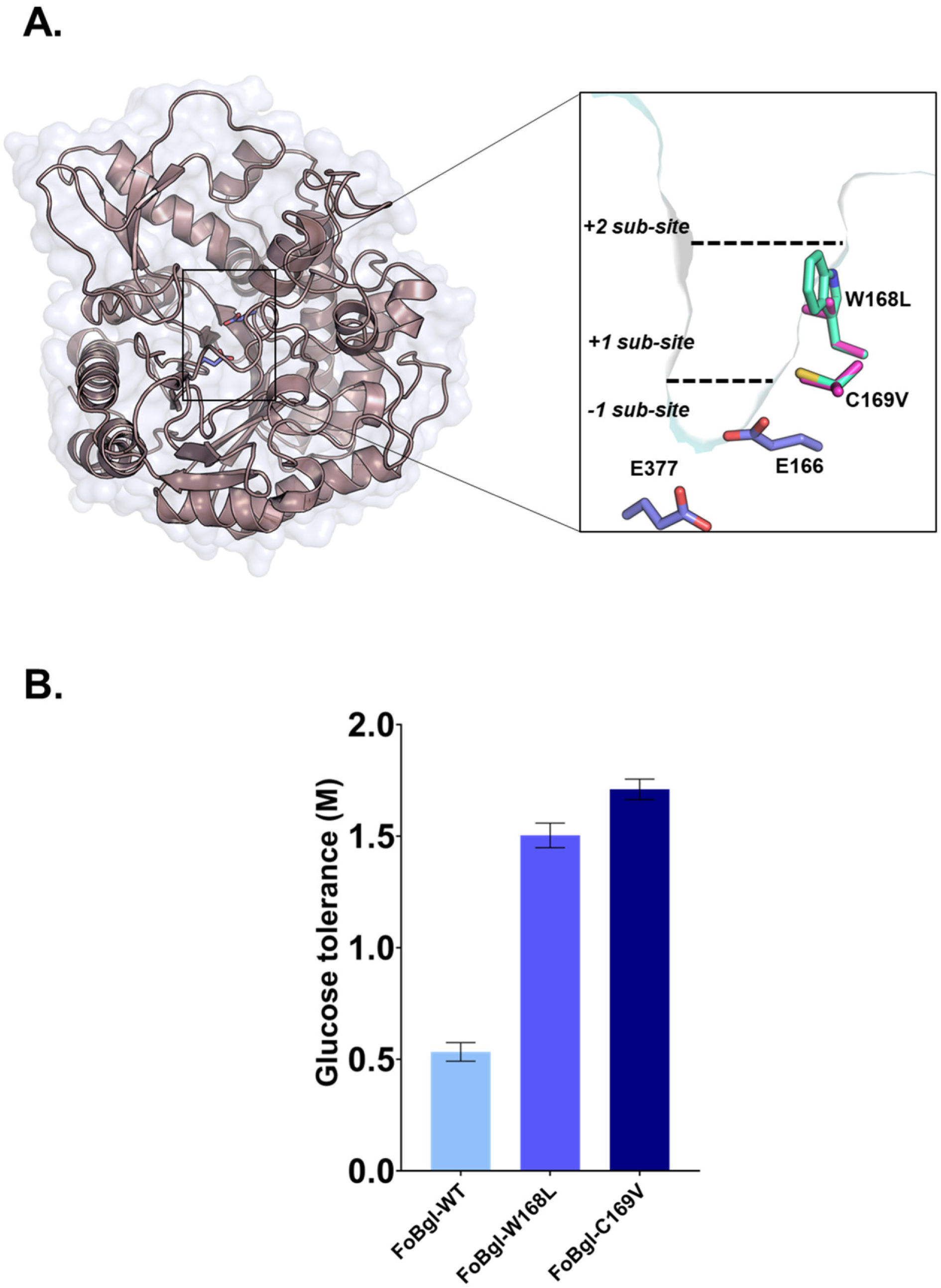
Mapping the residues forming the +1 sub-site of glucose binding inside the catalytic crater of FoBgl. (A) The overall structure of UnBGl1 is shown in the surface representation. Inset: zoomed-in view of the catalytic crater with the catalytic glutamates and residues present at the +1-sub-site shown in stick representation. (B) Comparison of the glucose tolerance levels of FoBgl-WT with those of the FoBgl-C169V and FoBgl-W168L mutants. The error bars correspond to the standard errors of the triplicate measurements.

#### Biochemical properties of +1 sub-site mutants of FoBgl

Biochemical assays of p-NPG and cellobiose hydrolysis were performed for all the mutants. Glucose tolerance assays were performed by using p-NPG hydrolysis. Both the mutants showed optimal p-NPG hydrolysis at 40°C and a pH of 6.0. Further, it was seen that both the +1 sub-site mutants of FoBgl showed improved glucose tolerance, estimated by p-NPG hydrolysis assays in the manner described in previous sections. The glucose tolerance of the FoBgl-C169V mutant was around 1.7 M, almost a 3-fold increase from that of FoBgl-WT. Whereas the FoBgl-W168L mutant showed glucose tolerance of around 1.3 M (Figure 4B). This indicated that the +1 sub-site has an essential role in product binding, and preventing the product glucose from stabilising at the +1 sub-site can improve the glucose tolerance of GH1 enzymes.

However, it was surprising that both these mutants were inactive against the actual substrate, cellobiose. Even after incubating high concentrations of these mutants (∼10 mg/ml) with cellobiose, it was observed that the C169V and W168L variants of FoBgl did not hydrolyse cellobiose. This finding led us to conclude that these residues at the +1 sub-site of FoBgl play an important role in the binding and stabilisation of the substrate, and mutation of these residues cannot be carried out for the improvement of glucose tolerance, as it adversely affects the entry and binding of the original substrate, cellobiose.

#### Mutations at the +2 sub-site of sugar-binding

As indicated in the preceding section, the mutations at the +1 sub-site of sugar-binding in FoBgl (FoBgl-W168L, FoBgl-C169V mutants) improve the glucose tolerance of the enzyme. However, such mutations also lead to total loss of cellobiose hydrolysis activity in FoBgl. This indicated that the +1 sub-site in FoBgl cannot be targeted for the generation of mutants to improve its glucose tolerance. Next, we targeted the +2 sub-site to assess its effects on the catalytic performance of FoBgl. The K256 residue close to the entry of the catalytic crater was mutated to hydrophobic amino acids. Both aromatic (tryptophan) and aliphatic (isoleucine and leucine) amino acid substitutions were carried out (Figures 5A-C). These mutants (FoBgl-K256W, FoBgl-K256I, and FoBgl-K256L) were expressed and purified in a similar manner as described previously. All mutants showed optimum cellobiose hydrolysis at 35-45°C (Figure 5D). The pH profile of these mutants was slightly distinct from FoBgl-WT. However, all these mutants except FoBgl-Y325F showed >50% activity over the pH range of 4.5-8.0 (Figure 5E). The FoBgl-K256W mutant showed an improved glucose tolerance of ∼ 0.9M compared to the FoBgl-WT (0.56 M) (Figure 5F). The improvement of the glucose tolerance can be attributed to the fact that the lysine residue in FoBgl-WT stabilised the glucose molecule by polar interactions at the +2 sub-site, leading to enzyme inhibition. However, the tryptophan residue in FoBgl-K256W mutant can interact with the carbon backbone of glucose molecules by stacking interactions, which has been reported in several GH family enzymes (44–46). The FoBgl-K256I and FoBgl-K256L mutants, thus generated, were expressed and purified in the same manner as FoBgl-WT. The assays revealed that the FoBgl-K256I mutant possessed far superior glucose tolerance compared to FoBgl-WT. It was observed that substitution of the K256 residue at the +2 sub-site of FoBgl caused a remarkable increase in glucose tolerance. The glucose tolerance of the FoBgl-K256I mutant was estimated to be around 1.2 M, two times greater than that of FoBgl-WT. FoBgl-K256L mutant also showed enhanced glucose tolerance of around 0.9 M. This shows that hydrophobic amino acids at the +2 sub-site prevent the stabilisation of product, improving the glucose tolerance of FoBgl.

**Figure 5:**
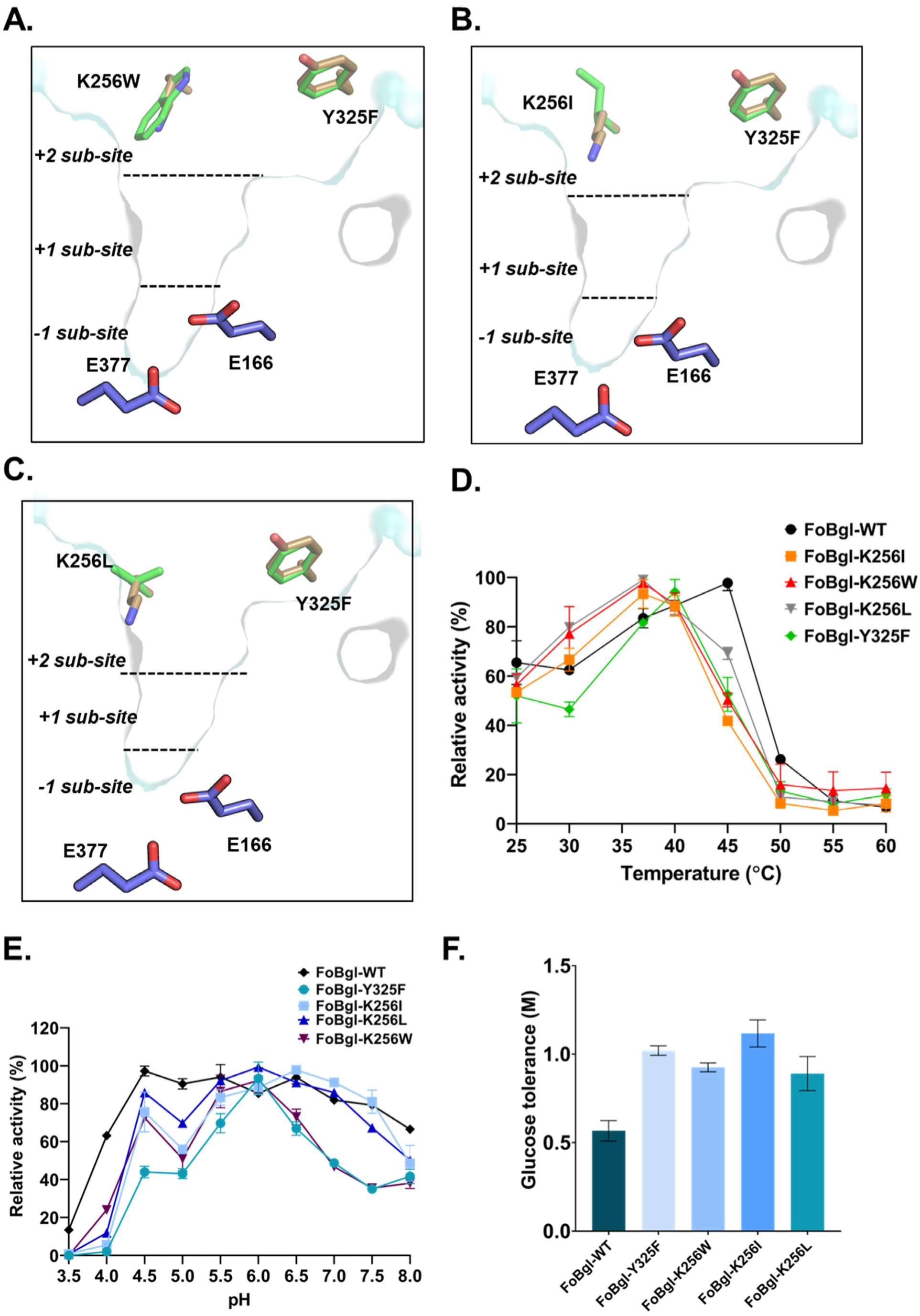
Mapping the residues forming the +2 sub-site of glucose binding inside the catalytic crater of FoBgl. (A, B, and C) The cross-section of the catalytic crater of FoBgl-WT with suggested mutations at the +2 sub-site shows the positions of K256W, K256I, K256L, and Y325F mutants. (D) Optimum temperature of cellobiose hydrolysis exhibited by each +2-sub-site single mutant. (E) Optimum pH of cellobiose hydrolysis exhibited by each +2-sub-site single mutant. (F) Effect of each mutation on the glucose tolerance of FoBgl. The error bars correspond to the standard errors of the triplicate measurements.

So far, we have successfully enhanced glucose tolerance by mutating the lysine residue present at the +2 sub-site of sugar-binding. On close examination of the structural model of FoBgl, we could observe a loop region guarding the entrance to the catalytic crater. This loop region forms the entrance to the catalytic crater of the enzyme. It was expected that substitutions at this loop would affect the interactions with the product molecule and might be a probable target for engineering for superior glucose tolerance. We observed the presence of a tyrosine residue at the 325^th^ position, capable of forming polar interactions with the product via its OH group, enabling product stabilisation, resulting in inhibition of the catalytic activity of FoBgl. The tyrosine residue was mutated to phenylalanine (FoBgl-Y325F) to eliminate the hydroxyl group, which was necessary to stabilise the sugar in its vicinity. This mutant was also expressed and purified, as mentioned previously. The assays revealed that FoBgl-Y325F showed glucose tolerance levels of ∼1.0 M (Figure 8C), a 400 mM increase compared to FoBgl-WT. Hence, we could establish the role of the tyrosine present in the elongated loop of FoBgl in interactions with the glucose moieties in its vicinity. Further, we also generated a double mutant, FoBgl-K256I-Y325F, combining the most glucose-tolerant variants at the +2 sub-site. This double mutant was expressed and purified similarly. The double mutants (FoBgl-K256I-Y325F and FoBgl-K256L-Y325F) exhibit optimal cellobiose hydrolysis at 35-40°C (Figure 6A). The pH profile indicated two distinct pH optima for both mutants, one at 4.5 and the other at 5.5 for FoBgl-K256W-Y325F, while the position of the second pH optima was observed to be at 6.5 for FoBgl-K256I-Y325F (Figure 6B) These mutants retained >50% activity over the broad pH range of 4.0-8.0, while the glucose tolerance assays showed a glucose tolerance of 1.4 M (Figure 6C). This shows that the lysine and tyrosine residues present at the +2 sub-site of the catalytic crater have roles in interacting and stabilising sugar moieties. Perturbation of these residues led to an overall enhancement of the glucose tolerance of FoBgl. It is important to note here that the cellobiose hydrolysis assays were also performed with each mutant. Unlike the mutation at the +1 sub-site, these mutants showed hydrolysis activity towards cellobiose.

**Figure 6:**
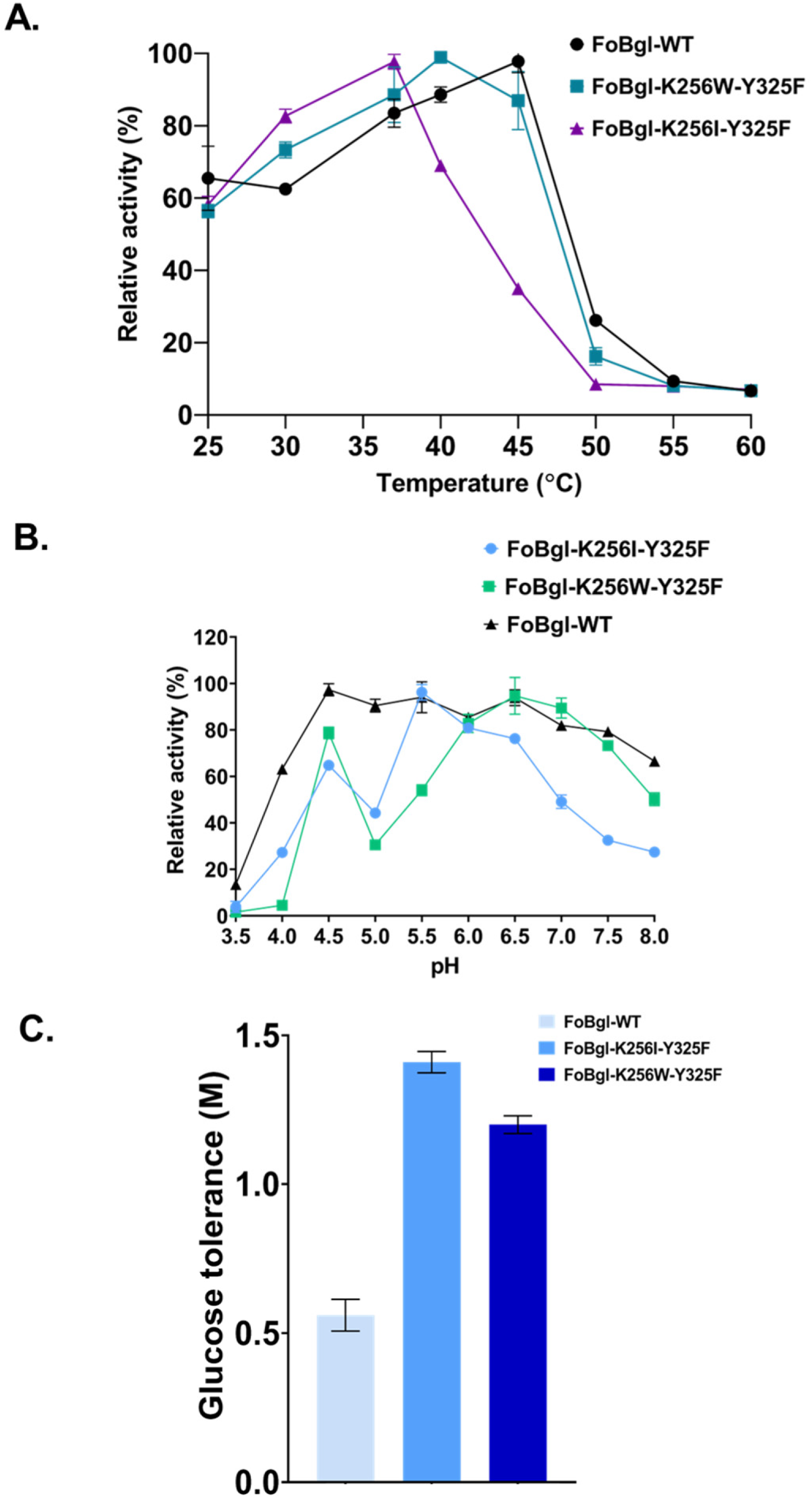
Biochemical effects of the double mutants targeting the +2 sub-site of glucose binding in FoBgl. (A) Optimum temperature of cellobiose hydrolysis (B) Optimum pH of cellobiose hydrolysis. (C) Glucose tolerance levels of the double mutants compared to FoBgl-WT. All error bars correspond to standard errors in readings taken in triplicate.

#### Hydrolysis assays with FoBgl mutants

##### Determination of kinetic properties

All mutants generated in the study adhered to the Michaelis-Menten enzyme kinetics model, displaying a hyperbolic curve of enzyme activity (or reaction velocity) against the substrate concentrations (Figure S3). The kinetic constants, *k_M_*, V_max_, *k_cat_*, and *k_cat_/k_M_* were determined to understand the catalytic efficiency of these mutants compared to FoBgl-WT. The values of the kinetic parameters of FoBgl mutants described in this study have been summarised in Table 1.

## Discussion

Exploring alternative energy sources is paramount to meet the growing population’s energy requirements. In this regard, the use of bioethanol has gained significant focus over recent years. The production of bioethanol from biomass requires the use of cellulase enzymes to break down cellulose into fermentable glucose. Here, we have characterised one β-glucosidase, the rate-limiting enzyme of the cellulase enzyme system. FoBgl-WT obtained from *Fusarium odoratissimum* has been characterized in detail in the present study. Highly pure recombinant FoBgl-WT was used for extensive biochemical characterisations using hydrolysis assays of p-NPG, the artificial substrate and its natural substrate cellobiose. The enzyme was found to carry out optimum hydrolysis of both p-NPG and cellobiose at a temperature of 40°C. However, the pH profiles for p-NPG and cellobiose hydrolysis were distinct. While the p-NPG hydrolysis profile showed a sharper peak corresponding to optimal activity at pH 6.0-6.5, no single peak was obtained for cellobiose hydrolysis. This is the first instance of the pH profiles of the same GH1 enzyme being so markedly different depending on the substrate used. FoBgl-WT retained more than 50% activity over a pH of 4.5-7.0. These biochemical properties make FoBgl-WT a promising candidate for industrial cellulose saccharification in second-generation bioethanol production. The industrial SSF reactors operate at 30-40°C and a pH of 5.0-6.0, which are also the optimal conditions for FoBgl-WT hydrolysis activity on cellobiose (47).

The structural model of the enzyme was generated in the absence of an experimentally determined structure. AlphaFold and SWISS-MODEL generated almost identical structures for FoBgl-WT, with the core structural fold identical to that of GH1 family enzymes. The model was further subjected to molecular dynamics (MD) simulation to assess stability. The RMSD and radius of gyration of the FoBgl-WT model remained stable throughout the 500 ns duration of the simulation process, indicating high quality and stability of the structure. The RMSF profile showed relatively higher fluctuations in residues present in the inherently flexible loop regions of the enzyme. In contrast, the rest of the residues, organised into proper secondary structures, exhibited very low fluctuations. The MD simulation approach was also used to understand the molecular details that determine the functionality of GH1 enzymes in general. Studies have reported that the GH1 enzymes exhibit a retaining mechanism of hydrolysis (48, 49), which is indicated by the presence of the two catalytic glutamates at a distance of ∼5Å. The distance between the CD of the E166 and E377 residues, which were presumed to be the catalytic residues, based on sequence and structural alignments with other well-characterised GH1 enzymes, remained ∼5.5Å throughout the 500 ns duration. Further, it is mentioned in the published literature that out of the two catalytic residues, the C-terminal glutamate acts as the catalytic nucleophile. In contrast, the N-terminal residue is the general acid/base in the catalytic process (50). Our simulation studies show the presence of a stable salt bridge interaction of E377 with R74, which might be the driving force behind maintaining this glutamate in a deprotonated state with an overall negatively charged carboxylate group. This deprotonated glutamate can then bring about a nucleophilic attack on the anomeric carbon of the substrate molecule. The R74 residue is conserved among all well-characterised GH1 enzymes in the literature. This strongly indicates the role of this conserved arginine in maintaining the glutamate, which serves as the catalytic nucleophile in a deprotonated state by charge-mediated interactions. The other glutamate, E166, lies within the conserved TXNEP motif, which is presumed to incorporate the catalytic residue serving as a general acid/base in the hydrolysis reactions. The principal role of this residue is to extract the proton from nearby water and generate activated water species, which brings about the second nucleophilic attack on the glucosylated enzyme intermediate to cause its breakdown and release the sugar product. For this step to be feasible, there must be an adequate number of water molecules which can interact with the E166 residue. Analysis of our MD simulation trajectories reveals the presence of multiple water molecules within the hydrogen bonding distance of the E166 residue. This indicates that this residue can readily abstract a proton from one of these water molecules and participate in the catalytic process as a general acid/base residue. Our simulation studies clearly indicate that the generated FoBgl-WT model as a stable three-dimensional structure. The MD simulation data and further structural analysis revealed important molecular details regarding the active site residues and their microenvironments, significantly contributing to maintaining this enzyme’s activity over a wide pH range. The finding of this study is expected to hold true for all GH1 enzymes with similar structural and functional attributes owing to conservation of the catalytic and substrate binding pocket residues.

Sequence alignments and structural comparisons of FoBgl-WT with UnBGl1, a glucose-tolerant GH1 family β-glucosidase obtained from the soil metagenomic library, were carried out. Our group has extensively characterised UnBGl1 both biochemically and structurally. UnBGl1 crystals diffract to atomic resolution (∼1 Å), and multiple crystal structures of UnBGl1 have been solved in apo-, substrate-, and product-bound forms. The glucose-bound crystal structures of UnBGl1 have shown that three glucose molecules are bound at specific sites within the catalytic crater (44). These conserved sites have been assigned as -1, +1, and +2 sub-sites for glucose binding. On an analysis of the structures of UnBGl1, it was observed that the polar residues lining the catalytic crater are crucial in stabilising the glucose molecules and, hence, inhibiting enzyme activity at higher glucose concentrations.

Structural studies on UnBGl1 and its structure-based rational engineering have demonstrated that residues lining the catalytic crater of the enzyme are essential for modulating the glucose tolerance. Mutation of a catalytic cysteine residue at the 188^th^ position to valine (C188V) resulted in a ∼2.5-fold enhancement of glucose tolerance in UnBGl1. This biochemical finding was further corroborated by the crystal structure of the C188V mutant of UnBGl1, which was solved in the glucose-bound state. The structure revealed the absence of glucose at the +1 sub-site of UnBGl1 C188V mutant. Similarly, the residues at the +2 sub-site were observed, which showed that a histidine residue at the 261^st^ position also stabilises another bound glucose at the entrance of the catalytic crater of UnBGl1. Mutation of this histidine residue to tryptophan (H261W) improved the glucose tolerance by ∼1.5-fold compared to wild-type UnBGl1 (44). Another GH1 β-glucosidase from the thermophilic bacterium *Acetivibrio thermocellus* (AtGH1) has been well characterised previously by our group (29). It was observed that substituting glycine (G168) with tryptophan (G168W) resulted in a ∼2.5-fold increase in glucose tolerance in AtGH1. This glycine residue is part of the +1 sub-site of glucose binding, as revealed by structural comparisons with UnBGl1. Mutation of glycine to tryptophan (AtGH1-G168W) constricted the catalytic crater due to the bulky side chain of the tryptophan residue that does not allow glucose to stabilise at that site. Guided by these previous studies on GH1 family enzymes performed in our group (29,44), we aimed to investigate the roles of the residues in +1 and +2 sub-sites in determining the glucose tolerance in FoBgl, as it exhibits high activity in a broad pH range. From the biochemical characterisations of FoBgl-WT by carrying out hydrolysis assays of different sugars, it was noted that FoBgl-WT can be used as a candidate for the saccharification reactions in SSF reactors. However, there was one major drawback in its industrial applicability. The glucose tolerance of FoBgl-WT was ∼500 mM glucose, meaning that the hydrolysis reaction gets inhibited by 50% at ∼500 mM product concentrations. The industrial reactors require the glucose tolerance of the enzymes to be ∼ 1M for their use. As a result, a structure-based rational engineering approach was undertaken to improve the glucose tolerance of the already pH-tolerant fungal GH1 β-glucosidase FoBgl. As mentioned previously, the core structural fold of FoBgl-WT was identical to that of the other characterised GH1 β-glucosidases. Structural alignments of FoBgl-WT with UnBGl1 revealed acceptable overall superposition, and the coordinates of the bound glucose in the UnBGl1 crystal structure could be used to map the probable glucose binding sites in FoBgl. The -1, +1, and +2 sub-sites of glucose binding were mapped in the catalytic crater of FoBgl based on the coordinates of the bound glucose molecules and the residues which interact with the glucose molecules were determined. The amino acids at the -1 sub-site were excluded from mutational studies as it was already known that this sub-site is essential for substrate binding. Any alteration at this site might result in loss of catalytic activity. Hence, the residues at the +1 sub-site were initially targeted for site-specific mutations. The cysteine at the +1 sub-site was conserved for both FoBgl-WT and UnBGl1. This C169 residue (numbered based on occurrence in FoBgl-WT) was mutated to valine, and we observed a tremendous increase in glucose tolerance from ∼500 mM to ∼1.7 M. However, the FoBgl-C169V mutant was inactive against cellobiose and only displayed p-NPG hydrolysis behaviour. The tryptophan at the 168^th^ position was also mutated to leucine (FoBgl-W168L) to mimic the exact residue present in UnBGl1. However, it was once again observed that the FoBgl-W168L mutant was inactive against cellobiose. This led us to conclude that, unlike UnBGl1, mutations at the +1 sub-site result in total loss of enzyme activity. Interestingly, glycine and valine are the residues at equivalent positions in AtGH1, another GH1 β-glucosidase. Despite bearing valine at the +1 sub-site of glucose binding, AtGH1 retains the cellobiose hydrolysis property. It has been observed that both UnBGl1 and AtGH1 β-glucosidases retain cellobiose-hydrolysing activity despite mutations at the +1 sub-site. Thus, the loss of cellobiose hydrolysis behaviour after mutations at the +1 sub-site is a unique observation in the case of FoBgl. Our biochemical assays performed using the mutants at the +1 sub-site of glucose binding suggest that every GH1 β-glucosidase has unique molecular determinants of substrate binding and hydrolysis kinetics. The +1 sub-site is crucial for substrate binding within the active site of the FoBgl. The fact that the +1 sub-site mutants show a total loss of cellobiose hydrolysis activity while retaining the p-NPG hydrolysis activity indicates that the natural and artificial substrates of this enzyme bind and interact inside the catalytic crater in two distinct fashions. This would also indicate that the biochemical characterisation of GH1 family β-glucosidases using p-NPG as a substrate for hydrolysis might not provide accurate mechanistic insights into the behaviour of these enzymes.

To enhance the glucose tolerance of FoBgl, we targeted the +2 sub-site of glucose binding as it was far from the catalytic crater, and the loss of activity on mutations at this site seemed less likely. The +2 sub-site of FoBgl-WT was observed to have a lysine residue in the 256^th^ position at a homologous site to the H261 residue in UnBGl1. We mutated K256 to tryptophan (FoBgl-K256W) to prevent glucose stabilisation via polar interactions. The result was encouraging as the glucose tolerance of this mutant was >1M, which is the industrial criterion for using these enzymes in fermenters. The kinetic properties of cellobiose hydrolysis were also similar to those of the wild-type enzyme. To further investigate the type of residues that might influence product stabilisation at the active site, we substituted the K256 residue with aliphatic amino acids such as leucine and isoleucine. For both these cases, we observed an increase in glucose tolerance levels compared to FoBgl-WT. However, the effect of such substitutions on the kinetic properties of the enzyme was quite interesting to note. Substitution of the lysine residue at the +2 sub-site to tryptophan results in an almost 2-fold increase in the catalytic efficiency of the enzyme. This might be attributed to better stabilisation of the substrate cellobiose at the entrance of the active site. The aromatic side chain of tryptophan also allows proper substrate orientation by providing stacking interactions with the carbon skeleton of the sugar moieties. This was further validated by substituting lysine with isoleucine and leucine at the same position. As expected, the glucose tolerance of the FoBgl-K256I mutant was even higher than that of FoBgl-K256W because the aliphatic side chain does not allow glucose to stabilise at the +2 sub-site. But it was interesting to note that there was no change in kinetic parameters and catalytic efficiency of FoBgl-K256I compared to that of the wild-type enzyme. The FoBgl-K256L mutant also showed improved glucose tolerance. The glucose tolerance of FoBgl-K256L was lower than that of the other two +2 sub-site variants. This is probably due to the smaller size of the leucine residue compared to the bulkier side chains of isoleucine and tryptophan residues.

The sequence and structure-based comparisons of FoBgl with other GH1 enzymes also revealed regions of insertions in the FoBgl sequence, resulting in longer loops in the 3D structure than in other GH1 enzymes. One such loop region borders the active site entrance and bears a tyrosine residue at position 325, which can stabilise the glucose produced after the hydrolysis of cellobiose via its OH group. We mutated this residue to phenylalanine (FoBgl-Y325F) and observed that the mutant showed ∼1M glucose tolerance, which was 2-fold higher than FoBgl-WT. Interestingly, the kinetic properties changed drastically by this substitution. The catalytic efficiency is reduced by ∼5-fold compared to FoBgl-WT. We also investigated the effect of combining the two +2 sub-site substitutions on the functional properties of FoBgl. In this regard, two double mutants, FoBgl-K256I-Y325F and FoBgl-K256W-Y325F, were generated. FoBg-K256I-Y325F showed the highest glucose tolerance (∼1.5 M) levels among all mutants generated in this study. A similar trend was also observed for the FoBgl-K256W-Y325F variant, which showed a glucose tolerance of ∼1.2 M. Interestingly, the catalytic efficiency and other kinetic properties were drastically affected by these substitutions, something we had also observed for the FoBgl-Y325F single mutant as well. This might indicate that substitutions at the 325th position had detrimental effects on FoBgl’s overall performance. Altercations at this position might destabilise the substrate, leading to reduced enzyme activity and turnover numbers. Although the mutations involving both residues at the +2 sub-site in FoBgl result in the highest glucose tolerance, a bargain exists with the catalytic efficiency of the mutant enzyme.

## Conclusions

This present study discusses the structural and functional characterisations of a GH1 β-glucosidase from the fungus *F. odoratissimum*. We have recombinantly expressed and obtained a highly pure form of the enzyme FoBgl from bacterial expression systems for the first time. Biochemical assays of cellobiose hydrolysis revealed FoBgl to be an extremely pH-tolerant enzyme, making it a promising candidate for the industrial saccharification processes. Structure-based rational engineering has been carried out to enhance the glucose tolerance of this enzyme. Mutations at the +2 sub-site of glucose binding resulted in a ∼2.5-fold improvement in glucose tolerance. We have successfully generated multiple mutants of FoBgl by structure-guided rational engineering with improved glucose tolerance; one of the mutants even shows an improvement in the catalytic efficiency.

Our study provides valuable insights into the molecular determinants of the kinetics of a GH1 enzyme. In this study, we report the detrimental effects of synonymous substitution at a site far away from the catalytic centre of the enzyme on the catalytic efficiency. This would act as an important guide for further rational engineering efforts for improvements in the properties of other enzymes belonging to the same family.

