## Supplementary Information for "Probing the role of residues lining the active site in the generation of glucose-tolerant variants of a fungal GH1 enzyme"

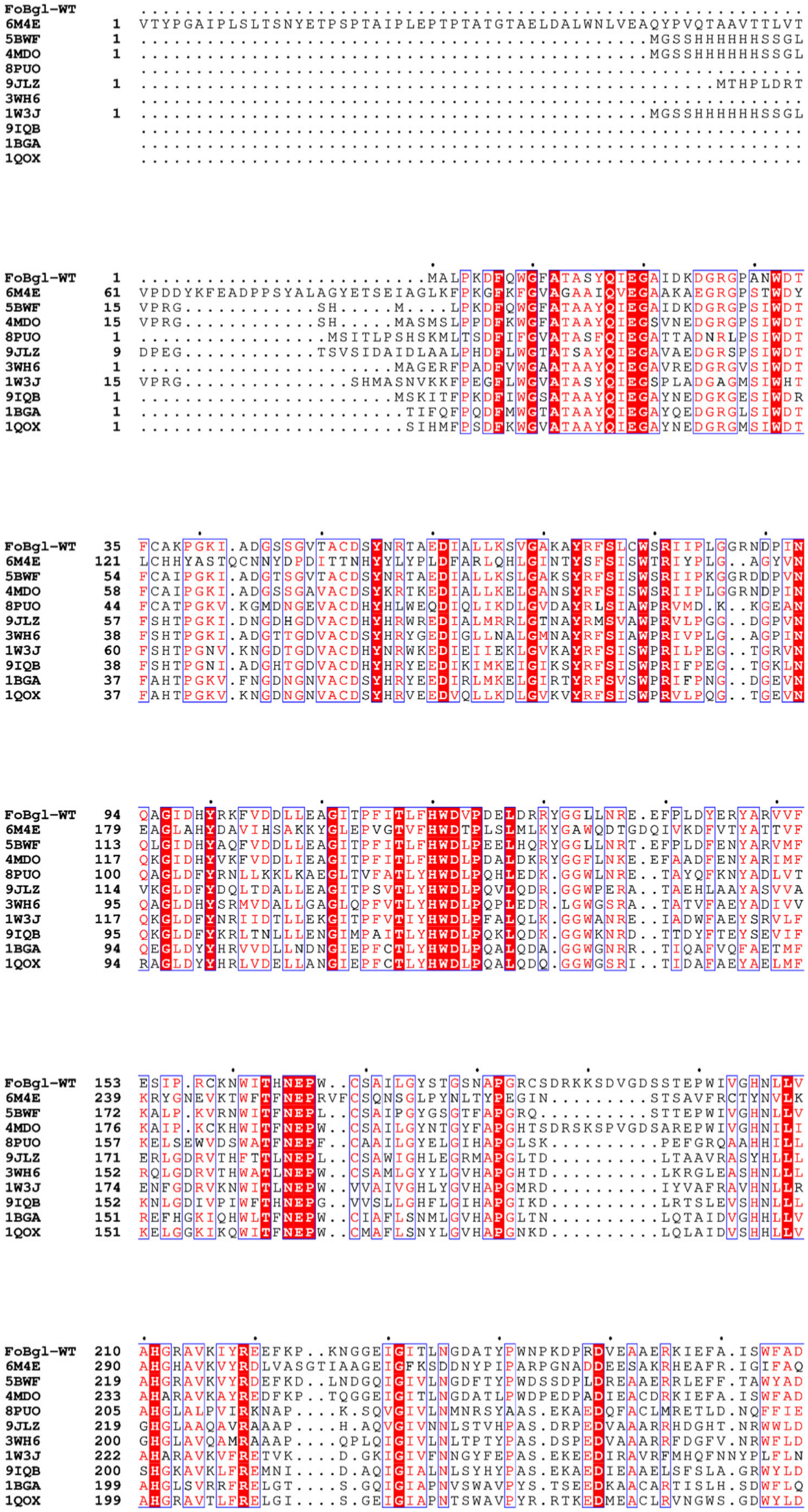

### Figure S1

```
FoBgl-WT   267   PIYF.GDYPASMRAQLGD..RLPTFTPEEKALVLGSNDFYGMNHYTANYVKHREGEAAPE
6M4E       349   PVYGNGDYPDVVKETVGD..MLPALTDEDKGYIKGSGDIFAIDGYRTDISHAALNG....
5BWF       276   PIYL.GDYPASMRKQLGD..RLPEFTPEEKAFVLGSNDFYGMNHYTSNYIRHRTSPATAD
4MDO       290   PIYF.GKYPDSMRKQLGD..RLPEFTPEEVALVKGSNDFYGMNHYTANYIKHKTGVPPED
8PUO       258   PLMK.GQYPQLLKTVAPQ..YLPTVLPGDMDIISQPIDFLGMNFYTCNHNAYDADDMFKN
9JLZ       272   PLHG.RGFPADMREVYGV..DLPE.RPGDLETIATPLDWLGLNYYFPAYIADDPDGPAPR
3WH6       254   PLAG.RGYPQDMLDYYGA..AAPQANPEDLTQIAAPLDWLGVNYYERMRAVDAPDASLPQ
1W3J       276   PIYR.GDYPELVLEFARE..YLPENYKDDMSEIQEKIDFVGLNYYSGHLVKFDPDAPAKV
9IQB       253   PVLK.GRYPENALKLYKKKGIELSFPEDDLKLISQPIDFIAFNNYSSEFIKYDPSSESGF
1BGA       252   PIYQ.GSYPQFLVDWFAEQGATVPIQDGDMDIIGEPIDMIGINYYSMSVNRFNPE..AGF
1QOX       252   PIYF.GEYPKFMLDWYENLGYKPPIVDGDMELIHQPIDFIGINYYTSSMNRYNPGEAGGM

FoBgl-WT   324   DYVGNLELHFWNHRGDCIGEET..............................QSTWLRPC
6M4E       403   .............IANCIRNQSDPNWPVCEEGSDPFAHVYPSGFAIGQSADPLSSWLVNS
5BWF       333   DTVGNVDVLFYNKEGQCIGPET..............................ESSWLRPC
4MDO       347   DFLGNLETLFYNKYGDCIGPET..............................QSFWLRPH
8PUO       315   V........KN....SQTVEYT..............................DIGW.EIA
9JLZ       328   AR......MVD....REGVPRT..............................GMGW.EID
3WH6       311   AQ......RLD....DPDLPHT..............................A.DR.EVY
1W3J       333   S........FV....ERDLPKT..............................AMGW.EIV
9IQB       312   SP.....ANSI....LEKFEKT..............................DMGW.IIY
1BGA       309   LQ.....SEEI....NMGLPVT..............................DIGW.PVE
1QOX       311   LS.....SEAI....SMGAPKT..............................DIGW.EIY

FoBgl-WT   354   ALGFRDLLVWISKRYGF.PRIYVTENGTSIKGENDMP.REKILQDDFRVKYYDDYVRAMA
6M4E       450   APFIRDQLKFLTQTYPAKGGIYFSEFGWAEDAEYDRQLLYQITWDGLRTQYLTDYLSQLL
5BWF       363   PAGFRDFLVWISKRYNY.PKIYVTENGTSLKGENDLP.KEKILEDDFRVNYYNEYIRAMF
4MDO       377   AQGFRDLLNWLSKRYGY.PKIYVTENGTSLKGENDMP.LEQVLEDDFRVKYFNDYVRAMA
8PUO       332   PQAFTELLVNLHKQYTL.PPIYITENGAACADQI....IDGEINDEQRVRYLDGHINAVN
9JLZ       347   ADGIETLLLRLTREYGA.RKLYVTENGSAFPDAV..R.PDGTVDDPERRDYLERHLAACA
3WH6       329   PEGLYDILLRLHNDYPF.RPLYITENGCALHDEI..A.EDGGIHDGQRQAFFEAHLAQLQ
1W3J       350   PEGIYWILKKVKEEYNP.PEVYITENGAAFDDVV..S.EDGRVHDQNRIDYLKAHIGQAW
9IQB       332   PEGLYDLLMLLDRDYGK.PNIVISENGAAFKDEI..G.SNGKIEDTKRIQYLKDYLTQAH
1BGA       329   SRGLYEVLHYL.QKYGN.IDIYITENGACINDEV....VNGKVQDDRRISYMQQHLVQVH
1QOX       331   AEGLYDLLRYTADKYGN.PTLYITENGACYNDGL..S.LDGRIHDQRRIDYLAMHLIQAS

FoBgl-WT   412   DASRLDGVDVHGYFAWSLLDNFEWAEGYETRFGVTYVDYEN.DQKRYPKKSAQHLKPLFD
6M4E       510   LAVHKDGINLRGALTWSFVDNWEWGLGMQQKFGFQFVNQSDPDLTRTFKLSAHAYAQFGR
5BWF       421   TAATLDGVNVKGYFAWSLMDNFEWADGYVTRFGVTYVDYEN.GQQRFPKKSAKSLKPLFD
4MDO       435   AAVAEDGCNVRGYLAWSLLDNFEWAEGYETRFGVTYVDYAN.DQKRYPKKSAKSLKPLFD
8PUO       387   H.AIESGVDIRGYFAWSLMDNFEWAEGYSKRFGLTYVDYQT..QERTIKRSGHAYRTLLN
9JLZ       403   S.AARRGAPLAGYFAWSLLDNFEWAYGYDKRFGLVHVDYAT..QTRTVKGSGHRYAEIIR
3WH6       385   R.ALAAGVPLKGYFAWSLLDNFEWAMGLSMRYGICYTNFET..LERRIKDSGYWLRDFIA
1W3J       406   K.AIQEGVPLKGYFVWSLLDNFEWAEGYSKRFGIVYVDYST..QKRIVKDSGYWYSNVVK
9IQB       388   R.AIQDGVNLKAYYLWSLLDNFEWAYGYNKRFGIVHVNFDT..LERKIKDSGYWYKEVIK
1BGA       383   R.TIHDGLHVKGYMAWSLLDNFEWAEGYNMRFGMIHVDFRT..QVRTPKESYYWYRNVVS
1QOX       387   R.AIEDGINLKGYMEWSLMDNFEWAEGYGMRFGLVHVDYDT..LVRTPKDSFYWYKGVIS

FoBgl-WT   471   SLIKQEEHAVNGNGVKAGQT
6M4E       570   NHL......HHHHHH.....
5BWF       480   ELIAKE..............
4MDO       494   SLIRKE..............
8PUO       444   NRKGLEHHHHHH........
9JLZ       460   AHRDGGQKAA..........
3WH6       442   GQRGKLAALEHHHHHH....
1W3J       463   NNGLED..............
9IQB       445   NNGF................
1BGA       440   NNWLETRR............
1QOX       444   RGWLDL..............
```

### Figure S1

**Figure S1: Sequence alignment of FoBgl-WT with structurally characterised GH1 enzymes**of different GH1 enzymes are mentioned as sequence identifiers. 6M4E: enzyme from *Hamamotoa singularis*, 5BWF: enzyme from *Trichoderma herzanium*, 4MDO: enzyme from *Humicola insolens*, 8PUO: enzyme from an Antarctic Marinomonas, 3WH6: enzyme Td2F2 from metagenome, 1W3J: enzyme from *Thermotoga maritima*, 1BGA: enzyme from *Bacillus polymyxa*, 1QOX: enzyme from *Bacillus circulans.* 9LJZ: enzyme from soil metagenome (UnBGl1), and 9IQB: enzyme from *Acetivibrio thermocellus*.

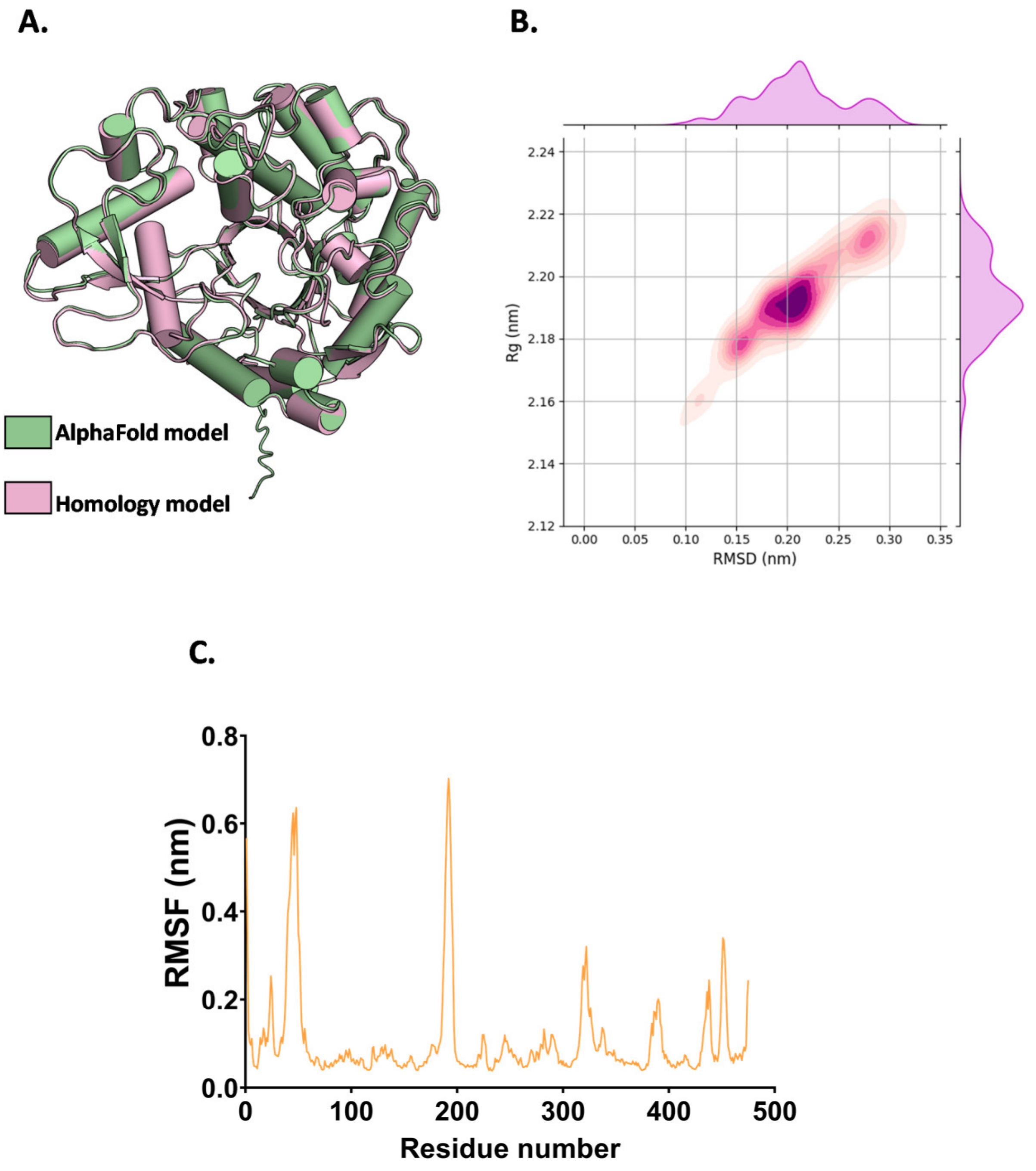


### Figure S2

**Figure S2: Analysis of the structure and molecular dynamics simulations to validate the stability of the generated FoBgl-WT model**. (A.) Structural superposition of the models of FoBgl-WT generated from AlphaFold (green) and SWISS MODEL (salmon) (B.) 2D contour plot comparing the RMSD with the $R_g$ (C.) RMSF plot showing fluctuations of the backbone of amino acid residues during 500 ns of MD simulation.

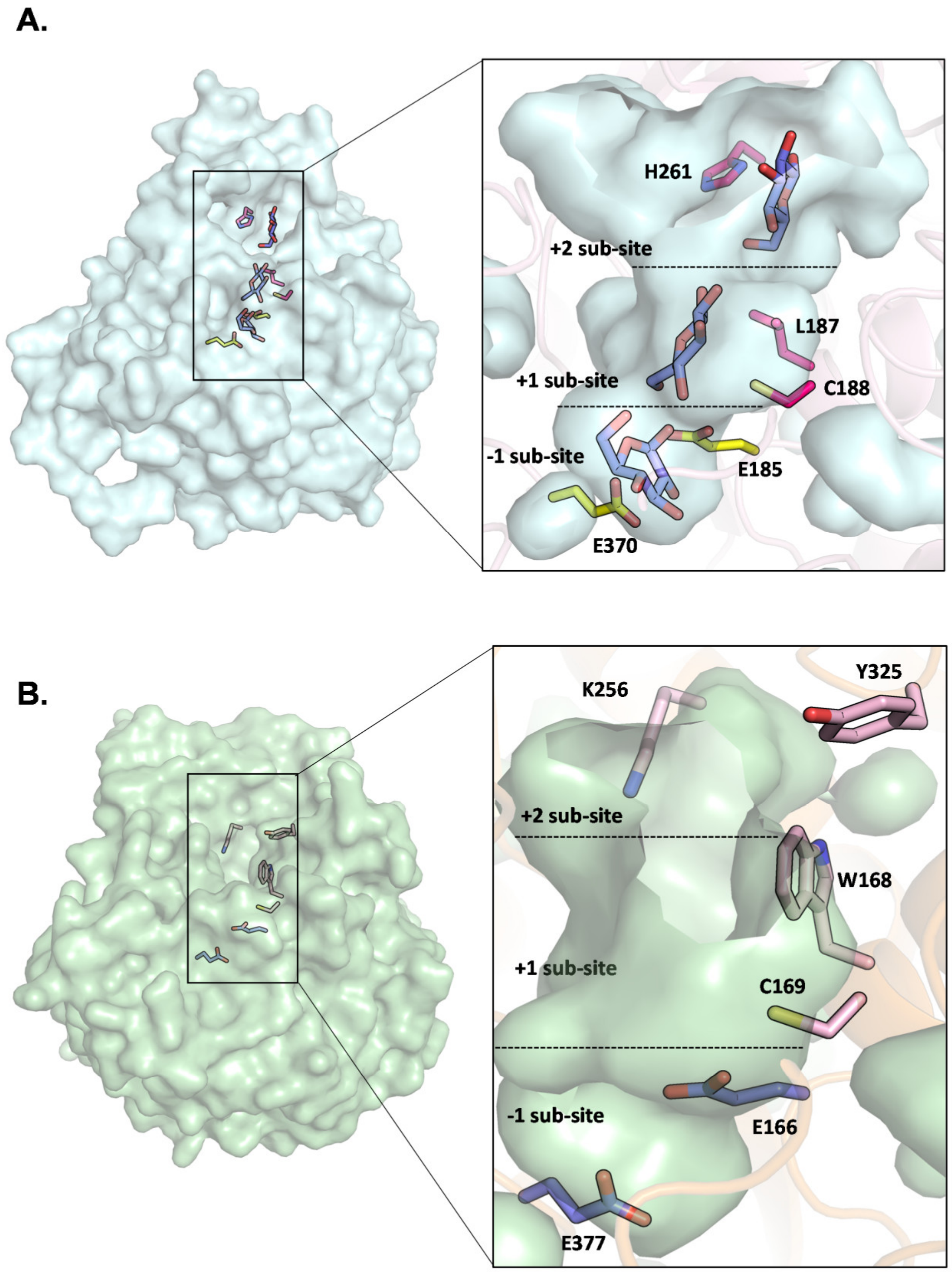


**Figure S3**

**Supplementary figure S3: Comparison of the active site architecture of UnBGl1 with that of FoBgl-WT.** (A.) Overall structure of UnBGl1 with the residues lining the catalytic crater and the bound glucose molecules shown in stick representation. Inset: Zoomed-in view of the catalytic crater with the catalytic glutamates depicted as yellow sticks and residues forming the +1 and +2 sub-sites of glucose binding shown as magenta sticks. The bound glucose molecules are shown as blue sticks. The -1, +1, and +2 sub-sites of glucose binding are indicated by dotted lines. (B.) Overall structure of FoBgl-WT with the residues lining the catalytic crater shown in stick representation. Inset: Zoomed-in view of the catalytic crater with the catalytic glutamates depicted as blue sticks and residues forming the +1 and +2 sub-sites of glucose binding shown as light pink sticks. The -1, +1, and +2 sub-sites of glucose binding are indicated by dotted lines.

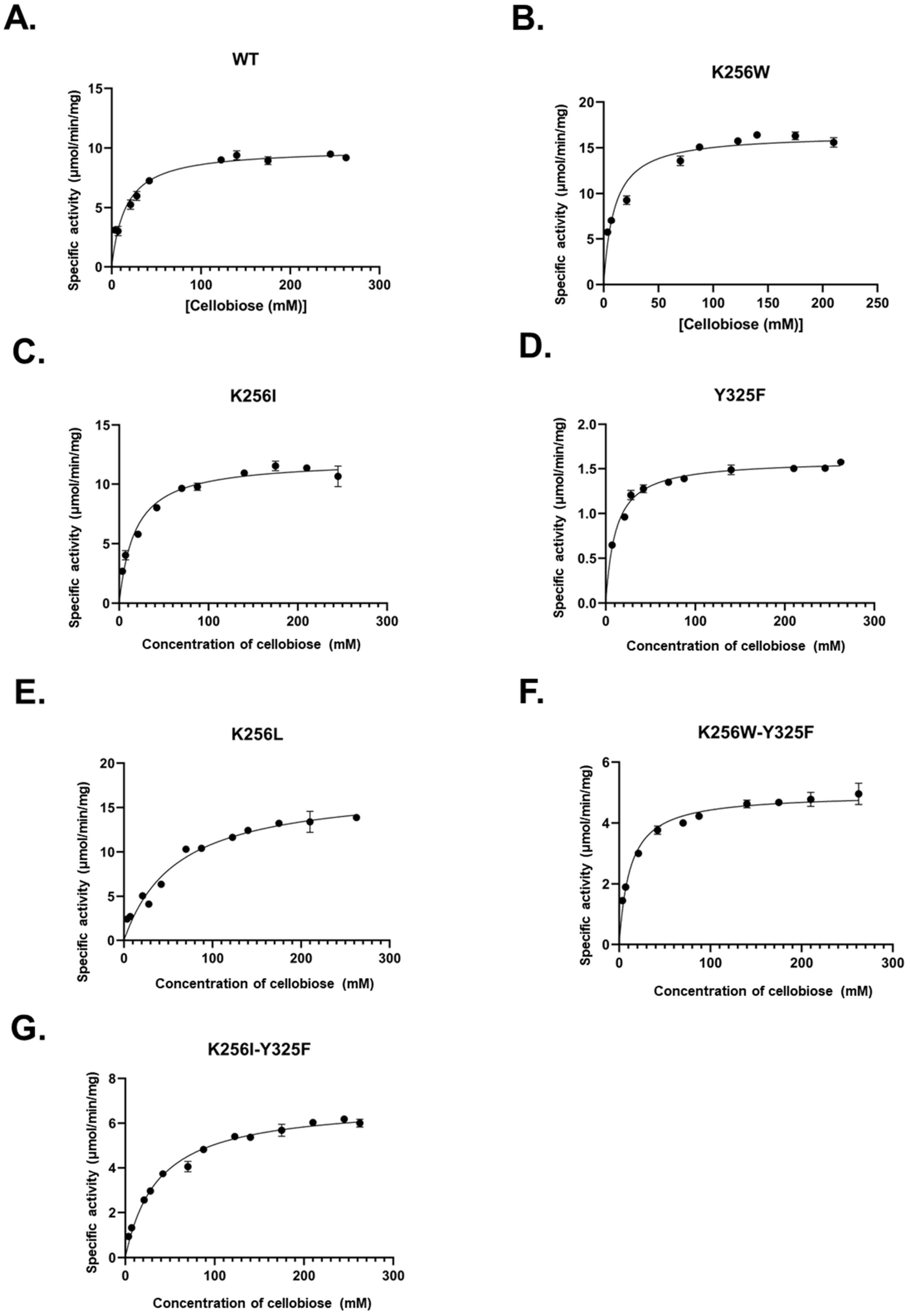
A.
WT
Specific activity (μmol/min/mg)
[Cellobiose (mM)]
B.
K256W
Specific activity (μmol/min/mg)
[Cellobiose (mM)]
C.
K256I
Specific activity (μmol/min/mg)
Concentration of cellobiose (mM)
D.
Y325F
Specific activity (μmol/min/mg)
Concentration of cellobiose (mM)
E.
K256L
Specific activity (μmol/min/mg)
Concentration of cellobiose (mM)
F.
K256W-Y325F
Specific activity (μmol/min/mg)
Concentration of cellobiose (mM)
G.
K256I-Y325F
Specific activity (μmol/min/mg)
Concentration of cellobiose (mM)

**Figure S4**

**Supplementary figure S4: Enzyme kinetics of FoBgl-WT and the +2 sub-site mutants performed using cellobiose as a substrate.** The error bars correspond to the standard errors in reading obtained in triplicate.

**Supplementary movie M1:** Residues lining the catalytic crater of FoBgl and forming the sites for binding and stabilisation of glucose molecules. The pink mesh shows the volume of the catalytic crater, and the catalytic E166 and E377 residues are shown as yellow sticks. Residues present along the catalytic crater that were selected for the site-directed mutagenesis studies are shown as light green sticks, while all other residues that are present lining the catalytic crater have been depicted as blue sticks.
